# Cellular Senescence Mediates Doxorubicin Chemotherapy-Induced Vascular Endothelial Dysfunction: Translational Evidence of Prevention with Senolytic Treatment

**DOI:** 10.1101/2025.04.19.649672

**Authors:** Ravinandan Venkatasubramanian, Sophia A. Mahoney, David A. Hutton, Nicholas S. VanDongen, Vienna E. Brunt, Nathan T. Greenberg, Abigail G. Longtine, Lukas Brandt, Andreas M. Beyer, Simon Melov, Judith Campisi, Matthew J. Rossman, Douglas R. Seals, Zachary S. Clayton

## Abstract

**Background:** Mechanisms underlying Doxorubicin (Doxo) chemotherapy-induced vascular endothelial dysfunction are incompletely understood.

**Objectives:** Determine the role of cellular senescence in mediating Doxo-induced vascular endothelial dysfunction and the influence of senolytic therapy as a therapeutic strategy to mitigate endothelial dysfunction with Doxo.

**Methods:** Endothelial function (carotid artery endothelium-dependent dilation [EDD] to increasing concentrations of acetylcholine) and associated mechanisms were assessed in young adult p16-3MR mice (which allow for genetic-based clearance of senescent cells with ganciclovir [GCV]) injected with Doxo and subsequently treated with GCV or ABT263 (senolytic). We also assessed the influence of Doxo and ABT263 *ex vivo* on EDD to increased flow in human arterioles.

**Results:** Lower peak EDD with Doxo (75±3% vs. control, 93±1%; P<0.05) was prevented with GCV (94±1%; P<0.05) and ABT263 (95±2%; P<0.05) treatment, which was mediated by preserved nitric oxide bioavailability and prevention of excess mitochondrial oxidative stress. In human arterioles, *ex vivo* Doxo exposure impaired peak EDD (Doxo, 32±10% vs. Control, 94±2%; P<0.05) which was prevented with concomitant incubation of Doxo with ABT263 (82±7%; P<0.05 vs. Doxo alone; P=0.63 vs. Control).

**Conclusion:** We provide translational evidence that cellular senescence contributes to Doxo-induced vascular endothelial dysfunction and that senolytics hold promise for preserving vascular endothelial function following Doxo exposure.

## INTRODUCTION

Cardiovascular diseases (CVD) and cancer are the two leading causes of mortality worldwide. There is estimated to be 2 million new cancer cases in the US this year^1^. Of these cases, ∼650,000 patients are expected to undergo chemotherapy treatment^1,2^. Although these therapies are effective at treating the cancer, they come with severe CV side effects. Chemotherapy-treated cancer survivors have a significantly higher risk of developing CVD later in life relative to age- and sex-matched healthy counterparts^3^. As a result, CVD are the leading cause of later morbidity and early mortality among cancer survivors treated with chemotherapy^1,3^. Anthracyclines, first-line chemotherapeutic agents for several common cancers, have toxic effects on the CV system^4,5^. Doxorubicin (Doxo) is the most widely used anthracycline^6,7^; thus, many patients treated with Doxo survive their cancers only to develop serious CV morbidity and die prematurely of CVD^1,6^,.

The great majority of CVD-related mortality following Doxo treatment stems from clinical atherosclerotic diseases^3^. The major antecedent of these CV disorders is endothelial dysfunction^8^. Doxo-related endothelial dysfunction, as shown by reduced endothelium-dependent dilation (EDD), is mediated primarily by reduced nitric oxide (NO) bioavailability^9^. This impairment in NO-mediated endothelial function is largely due to excessive mitochondrial superoxide-related oxidative stress, which reacts with NO, reducing its bioavailability^9^. However, the mechanistic event(s) governing excessive oxidative stress and endothelial dysfunction with Doxo are incompletely understood.

One likely mechanism contributing to Doxo-induced endothelial dysfunction is cellular senescence^5,10^, a state of cell cycle arrest coupled with the production of numerous pro-oxidative molecules, termed the senescence associated secretory phenotype (SASP)^11^. Physiological levels of cellular senescence are critical for many biological processes (e.g., cancer suppression; wound healing)^12^; however, excessive accumulation of senescent cells occurs in multiple tissues following Doxo treatment^5,10^. This accumulation can induce tissue dysfunction, at least in part through the SASP, and may be involved in select Doxo-induced pathologies. Thus, cellular senescence may contribute to a reduction in NO bioavailability, oxidative stress, and endothelial dysfunction with Doxo. However, the role of cellular senescence in mediating Doxo-induced endothelial dysfunction has not been investigated. Moreover, compounds that selectively clear senescent cells (termed senolytics) improve select indices of physiological dysfunction induced by Doxo chemotherapy^13–15^. Senolytic treatment following Doxo chemotherapy may hold promise for preventing endothelial dysfunction; however, the impact of senolytic treatment on Doxo-induced endothelial dysfunction has not yet been assessed.

Here, we performed three complementary studies. In study 1, we utilized the p16-3MR mouse model which allows for genetic-based clearance of excess senescent cells^16^, to establish the role of cellular senescence mediating endothelial dysfunction with Doxo. In study 2, we targeted cellular senescence with the pharmacological senolytic agent ABT263^15,17^ in mice to establish proof-of-principle efficacy of using senolytic therapy for preventing endothelial dysfunction with Doxo. In study 3, we translated our findings from study 2 to determine if ABT263 could prevent Doxo-induced endothelial dysfunction, *ex vivo*, in human arterioles. We used a variety of experimental approaches to elucidate the role of NO bioavailability and specifically excessive mitochondrial oxidative stress in regulating cellular senescence-mediated endothelial dysfunction with Doxo.

## MATERIAL AND METHODS

### Data Availability

Detailed description of the methods is available under **Supplementary Material**.

### Animals

All male and female mice were housed in a conventional facility on a 12-hour light/dark cycle, given ad libitum access to an irradiated, fixed, and open standard rodent chow (Inotiv/Envigo 7917) and drinking water. All mice were euthanized by cardiac exsanguination while maintained under anesthesia (inhaled isoflurane) 2 to 4 weeks following the completion of the intervention periods (allowing for 1 week of recovery). After cardiac exsanguination, the carotid arteries were excised under a stereoscope, dissected free of perivascular adipose tissue, and cannulated to fine glass canula within a pressure myograph system (DMT, Denmark) for assessment of vascular endothelial function. The thoracic aorta was excised, dissected free of surrounding tissue, sectioned, and stored (flash frozen for protein abundance by JESS capillary electrophoresis-based immunoblotting (Protein Simple, San Jose, CA) and immunofluorescence. Investigators were blinded to the treatment group for data collection and biochemical analyses. All animal protocols were approved by the University of Colorado Boulder Institutional Animal Care and Use Committee (protocol no. 2618) and complied with the National Institutes of Health Guide for the Care and Use of Laboratory Animals. Details on all procedures and antibodies are provided in the **Supplemental Material**.

#### Study 1: Genetic-based clearance of senescent cells with GCV in p16-3MR mice

p16-3MR mice were bred, weaned, and aged (to 4 months of age) in the animal care facility at the University of Colorado Boulder. At 4 months of age, male and female p16-3MR mice received either a single intraperitoneal injection of Sham (sterile saline) or Doxo (10 mg/kg in Sham). One week later, mice either received the vehicle (Veh; saline) or ganciclovir (GCV; 25 mg/kg/day in Veh) by intraperitoneal injection (IP) for 5 consecutive days, which is the standard approach for clearing senescent cells in this model, as we have described previously^17^. This equated to 4 groups/sex: Sham-Veh; Sham-GCV; Doxo-Veh; Doxo-GCV. For the primary outcomes of endothelial function, we studied n=8-15/sex/group.

#### Study 2: Senolytic-based clearance of senescent cells with ABT263 in mice

4-6 month old male and female p16-3MR mice received a single intraperitoneal injection of Sham (sterile saline) or Doxo (10 mg/kg in Sham). One week later, mice either received the vehicle (Veh; 10% EtOH, 30% PEG400, 60% Phosal 50 PG) or the senolytic ABT263 (50 mg/kg/day in Veh) by oral gavage on an intermittent one week on; two weeks off, one week on dosing paradigm, as we have previously described^18^. There were 4 groups/sex (Sham-Veh; Sham-ABT263; Doxo-Veh; Doxo-ABT263; n=10-12/group).

#### Study 3: *Ex vivo* human adipose tissue arteriole study – Influence of Doxo and ABT263

Human arterioles were isolated from fresh surgical discard or consented tissue in accordance with the approved protocols by the Institutional Review Board of the Medical College of Wisconsin and Froedtert Hospital. Arterioles from healthy subjects (≤1 risk factor for CVD) were received de-identified but with information about health status, co-morbidities and medication. Subject demographics are presented in **Supplemental Table 3**. Healthy human arterioles were incubated with 100 nM Doxo in PBS (as previously described^18^), 2.5 µM ABT263 in PBS or with a combination of both Doxo and ABT263 overnight (16-20h). These studies were approved in Medical College of Wisconsin IRB protocols PRO00001094 and PRO00028824.

#### Experimental procedures

##### Conduit artery endothelial function

Endothelium-dependent dilation (EDD) was assessed in response to increasing doses of acetylcholine (Sigma-Aldrich), while endothelium-independent dilation was evaluated using escalating concentrations of the exogenous nitric oxide donor sodium nitroprusside (Sigma-Aldrich) in isolated carotid arteries, as we have previously described^9,15,17^. Additional details, including the full list of pharmacological agents used in pharmaco-dissection experiments, can be found in the **Supplemental Materials**.

##### Endothelial function in human adipose tissue arterioles

Human arterioles (100-300 μm in diameter) were dissected from de-identified surgical discard tissue kept in HEPES buffer, (275 mM NaCl, 20 mM HEPES acid, 12 mM glucose, 7.99 mM KCl, 4.9 mM MgSO_4_, 3.2 mM CaCl_2_·2H_2_O, 2.35 mM KH_2_PO_4_ and 0.07 mM EDTA). Following the removal of connective tissue, arterioles were cannulated on glass pipettes tips with matched impedance in an organ chamber containing Krebs buffer (123 mM NaCl, 19 mM NaHCO3, 11 mM glucose, 4,7 mM KCl, 2.5 mM CaCl_2_,1.2 mM MgSO_4_, 1.2 mM KH_2_PO_4_ and 0.026 mM EDTA). The buffer was supplied with 5% CO to ensure that the pH of 7.4 was maintained throughout the experiment. Following the cannulation, arterioles were subsequently pressured to 60 mmHg at 37°C for 30min. Endothelin-1 (max. 20nM) was utilized to pre-constricted vessels to 30-50% of their pressurized diameter and then subjected to five different flow gradients, ranging from 5 – 100 cmH2O, which equal shear rates of 5 to 25 dynes /cm^2^. EID at the end of max flow was assessed using 100 µM Papaverine.

### Statistical Analyses

All details of all statistical analyses are available in the **Supplemental Material**. Unless otherwise noted, data are presented as mean ± SEM in the text, figures, and tables. Statistical significance was defined as α = 0.05. All analyses were conducted using Prism, version 10 (GraphPad Software, Inc., La Jolla, CA).

## RESULTS

### Animal characteristics

#### Study 1: p16-3MR Mouse Study with GCV Administration

To determine the causal role of cellular senescence in Doxo-induced vascular endothelial dysfunction, we utilized the p16-3MR mouse model, the reference-standard model for assessing the role of p16^INK4A^-associated cellular senescence in mediating physiological function^19^. This mouse model allows for selective genetic-based suppression of p16^INK4A^-positive senescent cells with the antiviral drug GCV^4,16,17^. Upon administration, GCV recognizes a herpes simplex thymidine kinase located on the p16-3MR transgene, which subsequently induces apoptosis of p16^INK4A^-positive senescent cells^12^. In the present study, young (4 months old) male and female p16-3MR mice received either a single intraperitoneal injection of Sham or Doxo and one week later were either treated with Veh or GCV for 5 consecutive days via IP injection **(Figure 1A)**. At the time of sacrifice, animals that received Doxo had significantly lower total body (*P* = 0.003), quadriceps (*P* = 0.002), and visceral adipose tissue (*P* < 0.0001) mass (relative to Sham-Veh control animals) which was not prevented by suppression of excess senescent cells with GCV. No characteristic differences were observed between Sham-Veh and Sham-GCV animals. We observed no group differences in carotid artery resting or maximal diameter. (**Table 1**).

**Figure 1.**
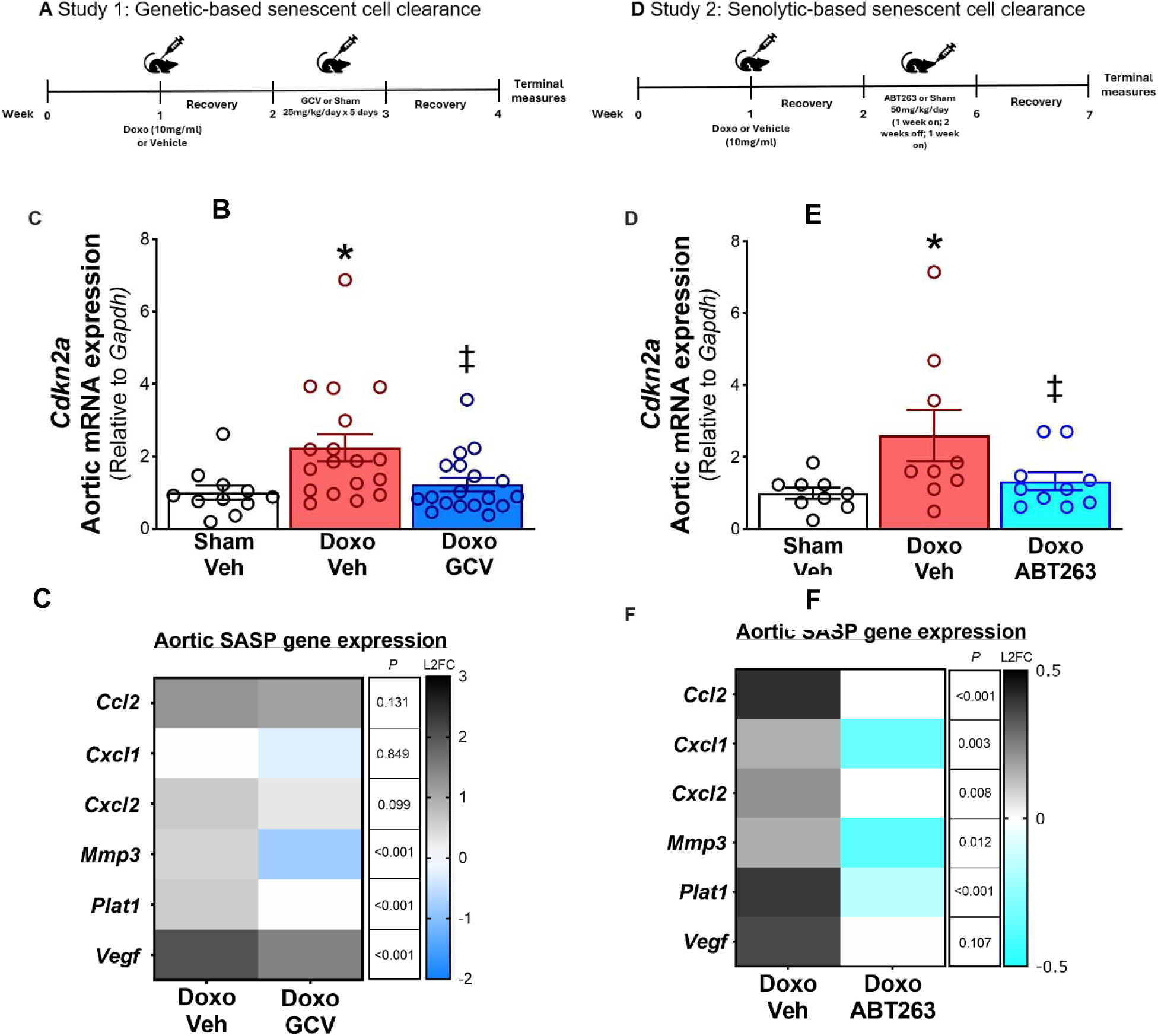
Doxorubicin (Doxo) administration increases aortic cellular senescence in young mice and senolytic treatment prevents it. Study paradigm of genetic-based clearance with ganciclovir (GCV) (Study 1) **(A)**. Senescent cell **(B)** and senescence-associated secretory phenotype (SASP) **(C)** marker expression for study 1. Study paradigm of senolytic-based clearance with ABT263 (Study 2) **(D)**. Senescent cell **(E)** and SASP **(F)** marker expression for study 2. All values are mean ± SEM. N=11-19/group in study 1, N=9-10/ group in study 2. *p<0.05 vs Sham-Vehicle (Veh), ‡p<0.05 vs Doxo-Veh.

**Table 1.** Characteristics of mice in study 1. Data are mean ± SEM. \**P*<0.05 vs. Sham-Vehicle (Veh).

|  | Sham<br>Veh | Sham<br>GCV | DOXO<br>Veh | DOXO<br>GCV |
| --- | --- | --- | --- | --- |
| ***n*** | 27<br>10 female<br>17 male | 20<br>9 female<br>11 male | 18<br>9 female<br>9 male | 26<br>14 female<br>12 male |
| **Body weight, g** | 30.0 ± 1.1 | 31.2 ± 1.3 | 25.3 ± 0.9* | 25.5 ± 0.6* |
| **Heart mass, mg** | 143.1 ± 3.5 | 143.2 ± 4.2 | 132.5 ± 5.7 | 132.9 ± 3.3 |
| **Left ventricle mass, mg** | 81.9 ± 3.9 | 80.1 ± 3.8 | 79.2 ± 4.9 | 78.7 ± 3.9 |
| **Quadriceps mass, mg** | 281.5 ± 9.9 | 278.1 ± 17.9 | 233.2 ± 9.8* | 242.3 ± 6.5* |
| **Visceral adipose mass, mg** | 787.5 ± 99.9 | 847.1 ± 127.7 | 203.9 ± 37.7* | 212.0 ± 31.9* |
| **Spleen mass, mg** | 75.9 ± 2.5 | 82.9 ± 2.8 | 77.6 ± 5.3 | 82.9 ± 3.7 |
| **Carotid artery** |  |  |  |  |
| **Resting diameter, μM** | 383 ± 8 | 397 ± 8 | 398 ± 7 | 391 ± 7 |
| **Maximal diameter, μM** | 446 ± 4 | 463 ± 7 | 468 ± 7 | 462 ± 7 |

#### Study 2: Senolytic (ABT263) Administration Mouse Study

We next aimed to determine the efficacy of targeting excess cellular senescence with a pharmacological senolytic to prevent Doxo-related vascular endothelial dysfunction. We selected ABT263 as it is a well-established synthetic pharmacological senolytic agent that we recently used to successfully target excess cellular senescence in old mice, resulting in improved vascular endothelial function^17^, and applied the same approach in the present study. Importantly, the effects of ABT263 on Doxo-induced vascular endothelial dysfunction had not yet been determined. The mechanism of action by which ABT263 elicits its senolytic properties is to lower the activity of BCL-2 (an anti-apoptotic pathway which is increased in senescent cells) and selectively induce apoptosis in senescent cells^14^. Here, we used young (4-month-old) male and female p16-3MR mice. One week after a single IP injection of Sham or Doxo we treated animals with either Veh or ABT263 (**Figure 1D**). At the time of sacrifice, animals that were administered with Doxo had lower visceral adipose tissue mass relative to Sham-administered animals (*P* = 0.041), which was not prevented by ABT263 treatment. No characteristic differences were observed between Sham-Veh and Sham-ABT263 animals. We observed no group differences in carotid artery resting or maximal diameter (**Supplemental Table 1**).

### Doxo administration results in elevated cellular senescence and SASP burden in arteries: prevention with genetic-based clearance of senescent cells and senolytic therapy

To establish whether cellular senescence and the SASP were greater in the vasculature following Doxo administration and if genetic-based clearance of senescent cells and senolytic therapy prevented greater senescence and SASP burden, we assessed a canonical biomarker of cellular senescence in mouse aorta lysates via RT-qPCR. We found that C*dkn2a* expression, the gene which encodes the p16^INK4A^ protein and the senescence marker implicated in vascular endothelial dysfunction^13,15,17^, was significantly higher in the Doxo-Veh group (vs. Sham-Veh) in studies 1 (2.24-fold higher, *P* = 0.019; **Figure 1B**) and 2 (3.23-fold, *P* = 0.035; **Figure 1E**). The Doxo-induced increase in *Cdkn2a* was prevented with genetic-based clearance of senescent cells in study 1 (Doxo-GCV vs. Doxo-Veh, *P* = 0.020; Dox-GCV vs. Sham-Veh, *P* = 0.438; **Figure 1B**) and senolytic treatment in study 2 (Doxo-ABT263 vs. Doxo-Veh, *P* = 0.042; Doxo-ABT263 vs. Sham-Veh, *P* = 0.164; **Figure 1E**). Considering ABT263 elicits its senolytic effects by reducing BCL2 activity, we quantified the BAX:BCL2 ratio, which is an established surrogate for BCL2 activity^14,20,21^. Relative to Sham-administered animals, the BAX:BCL2 ratio was 20% higher in aortic lysates from animals administered with Doxo (Doxo-Veh vs. Sham-Veh, *P* = 0.047), which was fully prevented by ABT263 treatment (Doxo-ABT263 vs. Doxo-Veh, *P* = 0.042; Doxo-ABT263 vs. Sham-Veh, *P* = 0.947; **Figure S1**). In addition to assessing biomarkers of senescent cell burden in arteries, we measured the expression of key SASP genes, including *Ccl2*, *Cxcl1*, *Cxcl2*, *Mmp3*, *Plat 1*, and *Vegf*^22,23^. We found in both studies 1 (**Figure 1C**) and 2 (**Figure 1F**) these genes were generally higher with Doxo (Doxo-Veh vs. Sham-Veh) and higher expression levels were prevented with genetic-based clearance of senescent cells with GCV (**Figure 1C**) and following senolytic therapy (**Figure 1F**). Overall, these results suggest that systemic Doxo administration results in elevated cellular senescence and SASP burden in the vasculature, which can be mitigated with systemic genetic-based clearance of excess senescent cells and systemic senolytic treatment.

### Cellular senescence underlies conduit artery endothelial dysfunction following Doxo administration: Prevention with senolytic treatment

#### Conduit Artery Endothelial Function

To determine whether Doxo administration causes conduit artery endothelial dysfunction as a result of excess cellular senescence, we assessed carotid artery EDD to increasing doses of ACh *ex vivo* in study 1. Peak EDD was 20% lower following Doxo administration (Doxo-Veh, 75 ± 3% vs. Sham-Veh, 94 ± 1%, *P* < 0.0001), which was fully prevented with genetic-based clearance of senescent cells (Doxo-GCV, 94 ± 1% vs. Doxo-Veh, *P* < 0.0001; Doxo-GCV vs. Sham-Veh, *P* = 0.99). Importantly, the effects of GCV were unique to Doxo-induced senescence, as evidenced by no influence of GCV on peak EDD in animals administered the Sham (Sham-GCV, 96 ± 1% vs. Sham-Veh, *P* = 0.94) (**Figure 2A**).

**Figure 2.**
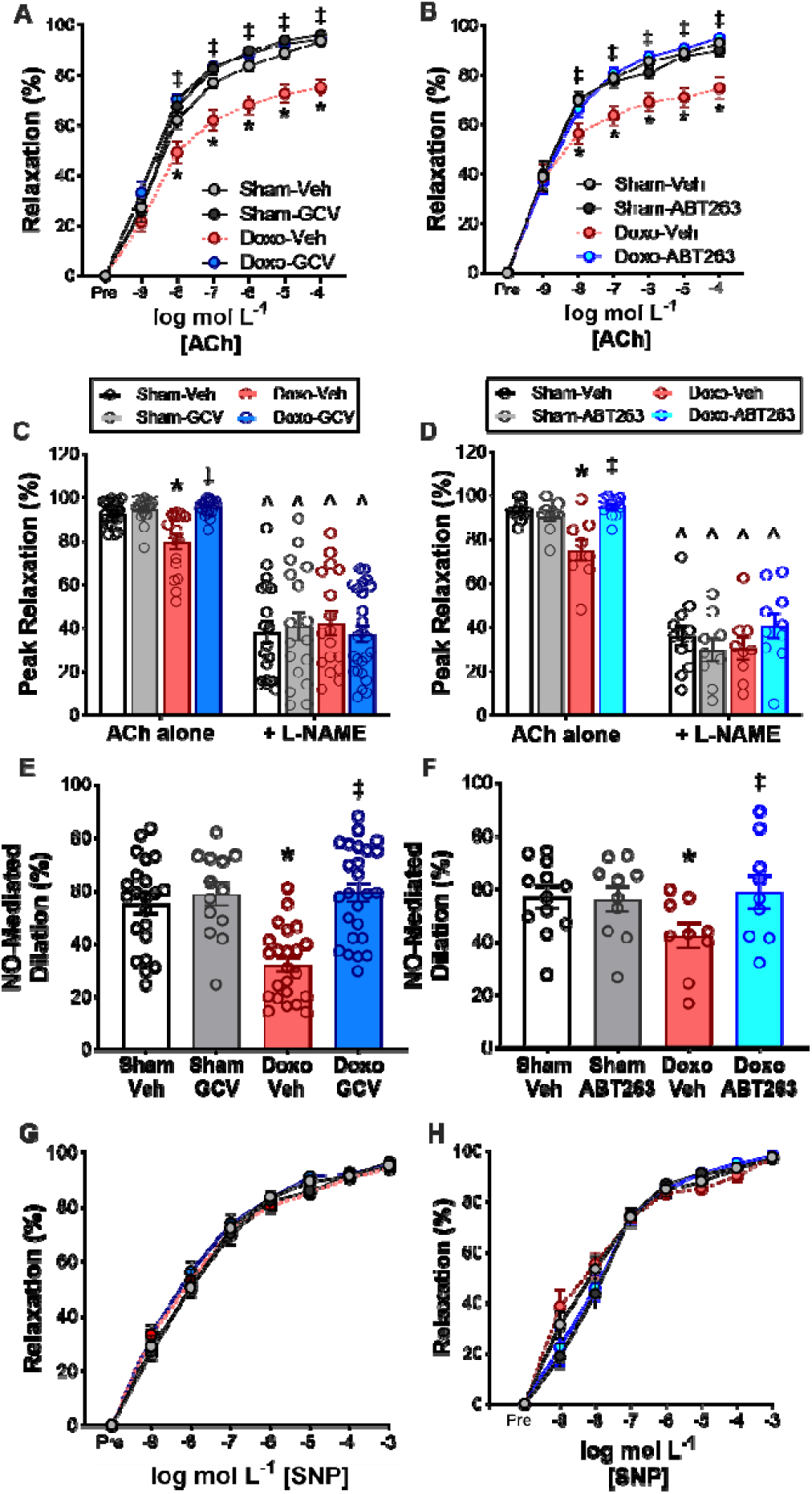
Doxo-induced cellular senescence impairs endothelial dysfunction by reducing nitric oxide (NO) bioavailability and not by influencing vascular smooth muscle sensitivity to NO, and senolytic treatment prevents these deleterious changes. Carotid artery endothelium-dependent dilation (EDD) to increasing doses of acetylcholine (ACh) in study 1 **(A)** and 2 **(B)**. NO-mediated EDD response to ACh in absence/presence of NO-synthase inhibitor, L-NAME in study 1 **(C)** and study 2 **(D)**. NO-mediated dilation calculated as peak EDD [–] EDD in presence of L-NAME in study 1 **(E)** and study 2 **(F)**. Endothelium independent dilation (EID) in response to NO-donor, sodium nitroprusside (SNP) in study 1 **(G)** and 2 **(H).** All values are mean ± SEM. N=13-25 in study 1, N=9-12 in study 2. *p<0.05 vs Sham-Veh, ‡p<0.05 vs Doxo-Veh. ^p<0.05, Ach alone vs Ach+MitoQ.

We next sought to determine if targeting excess cellular senescence with senolytic therapy could prevent Doxo-induced conduit artery endothelial dysfunction. To accomplish this, we again measured carotid artery EDD to increasing doses of ACh *ex vivo* in study 2. Peak EDD was 19% lower following Doxo administration (Doxo-Veh, 75 ± 4% vs. Sham-Veh, 93 ± 1%, *P* = 0.002), which was fully prevented with senolytic therapy (Doxo-ABT263, 95 ± 1% vs. Doxo-Veh, *P* < 0.0001; Doxo-ABT263 vs. Sham-Veh, *P* = 0.95). The influence of ABT263 was specific to Doxo-induced senescence, as evidenced by no influence of ABT263 on peak EDD in animals administered the Sham (Sham-ABT263, 90 ± 2% vs. Sham-Veh, *P* = 0.87) (**Figure 2B**).

Together, these results establish excess cellular senescence in the vasculature as an underlying mechanism of Doxo-induced conduit artery endothelial dysfunction and establish proof-of-concept efficacy for the use of senolytic therapy as a strategy for preventing Doxo-induced conduit artery endothelial dysfunction.

#### NO-Mediated Endothelial Function

We have previously shown that Doxo administration causes vascular endothelial dysfunction as a result of reduced NO bioavailability^9^. Thus, we sought to determine in study 1, whether a reduction in NO bioavailability with Doxo administration was mediated by excess cellular senescence. To accomplish this goal, we incubated carotid arteries with the endothelial NO synthase inhibitor L-NAME and measured EDD to increasing doses of ACh. L-NAME abolished group differences in peak EDD (**Figure 2C**), suggesting that differences in NO bioavailability underlie endothelial dysfunction with Doxo administration. Peak NO-mediated dilation (peak EDD with ACh alone [-] peak EDD with ACh plus L-NAME) was approximately 45% lower in Doxo-Veh mice compared to Sham-Veh mice (Sham-Veh, 55 ± 4% vs, Doxo-Veh, 30 ± 3%; *P* < 0.0001; **Figure 2E**), which was fully prevented by suppression of excess senescent cells with GCV (Doxo-GCV, 59 ± 4%; *P* < 0.0001 vs. Doxo-Veh; **Figure 2E**). The effects of GCV were unique to Doxo-induced senescence as evidenced by no differences in NO-mediated EDD between the Sham-Veh and Sham-GCV groups (Sham-GCV, 55 ± 5% vs. Sham-Veh, *P* = 0.99; **Figure 2E**).

Next, in study 2, we aimed to determine whether targeting excess cellular senescence with ABT263 treatment can preserve NO bioavailability following Doxo administration. Similar to study 1, L-NAME abolished group differences in peak EDD (**Figure 2D**). Peak NO-mediated EDD was 26% lower in Doxo-Veh mice compared to Sham-Veh mice (Sham-Veh, 57 ± 4%; Doxo-Veh, 42 ± 5%; *P* = 0.027. **Figure 2F**). The influence of ABT263 was specific to Doxo-induced senescence, as evidenced by no differences in NO-mediated EDD between the Sham-Veh and Sham-ABT263 groups (Sham-ABT263, 56 ± 5% vs Sham-Veh, *P* = 0.99; **Figure 2F**).

To determine if suppression of excess senescent cells following Doxo administration altered smooth muscle sensitivity to NO, we measured EID and found no group differences in study 1 (*P* = 0.12; **Figure 2G**) or study 2 (*P* = 0.47; **Figure 2H**). Together these results suggest that cellular senescence contributes to conduit artery endothelial dysfunction with Doxo administration by lowering NO bioavailability and not influencing vascular smooth muscle sensitivity to NO. Additionally, senolytic treatment following Doxo administration may be an effective therapy to preserve NO bioavailability.

#### Mitochondrial oxidative stress-mediated endothelial function

We have previously shown that Doxo administration results in excess superoxide-related oxidative stress in the vasculature, which directly contributes to Doxo-induced conduit artery endothelial dysfunction^9^. Thus, we first sought to determine if ABT263 treatment could prevent excessive oxidative stress in the vasculature following Doxo administration. Nitrotyrosine, a biomarker of oxidative protein damage, is produced when cellular superoxide reacts with NO to form peroxynitrite (ONOO-), which can then undergo chemical oxidation with protein tyrosine to form nitrotyrosine^24^. We found that higher nitrotyrosine levels with Doxo administration was mitigated by ABT263 senolytic treatment (Doxo-Veh, 876 ± 29 vs. Doxo-ABT263, 799 ± 53 Arbitrary Units [AU]; *P* = 0.036; **Figure S2A**).

After establishing that excessive vascular oxidative stress with Doxo was mitigated by senolytic treatment, we then sought to determine the role of mitochondria as a source of this excessive oxidative stress and whether it was regulated by cellular senescence. We have previously shown that excess mitochondrial reactive oxygen species (mitoROS)-related oxidative stress is implicated in Doxo-induced vascular endothelial dysfunction^8^. Therefore, in study 1 we first aimed to investigate the role of excess cellular senescence in mediating excessive mitoROS induced by Doxo administration. To accomplish this, we assessed mitoROS bioactivity in isolated aortic segments by electron paramagnetic spectroscopy, the reference standard experimental approach for measuring mitoROS bioactivity in biological tissues^25^. We found that the Doxo-Veh group had 2.3-fold higher aortic mitoROS bioactivity compared to the Sham-Veh group (Doxo-Veh, 7286 ± 1429 AU vs. Sham-Veh, 3116 ± 634 AU; *P* = 0.006; **Figure 3A**). Genetic-based clearance of excess senescent cells with GCV following Doxo administration prevented greater mitoROS bioactivity with Doxo (Doxo-GCV, 3272 ± 811 AU; *P* = 0.021 vs. Doxo-Veh; **Figure 3A**). The effects of GCV were unique to Doxo-induced senescence as evidenced by no differences in mitoROS bioactivity between the Sham-GCV (2782 ± 598 AU) and Sham-Veh groups (*P* = 0.71; **Figure 3A**). Similarly, in study 2, we found that Doxo-Veh mice had higher aortic mitoROS bioactivity relative to Sham-Veh animals (Doxo-Veh, 14296 ± 1238 AU vs. Sham-Veh, 9272 ± 1088 AU, *P* = 0.011; **Figure 3B**). Moreover, clearance of excess senescent cells with senolytic therapy prevented greater mitoROS bioactivity with Doxo (Doxo-ABT263, 9063 ± 1129 AU; *P* = 0.009 vs. Doxo-Veh; **Figure 3B**). The effects of ABT263 were unique to Doxo-induced senescence as evidenced by no differences in mitoROS bioactivity between the Sham-ABT263 (9622 ± 1655 AU) and Sham-Veh groups (*P* = 0.86; **Figure 3B**).

**Figure 3.**
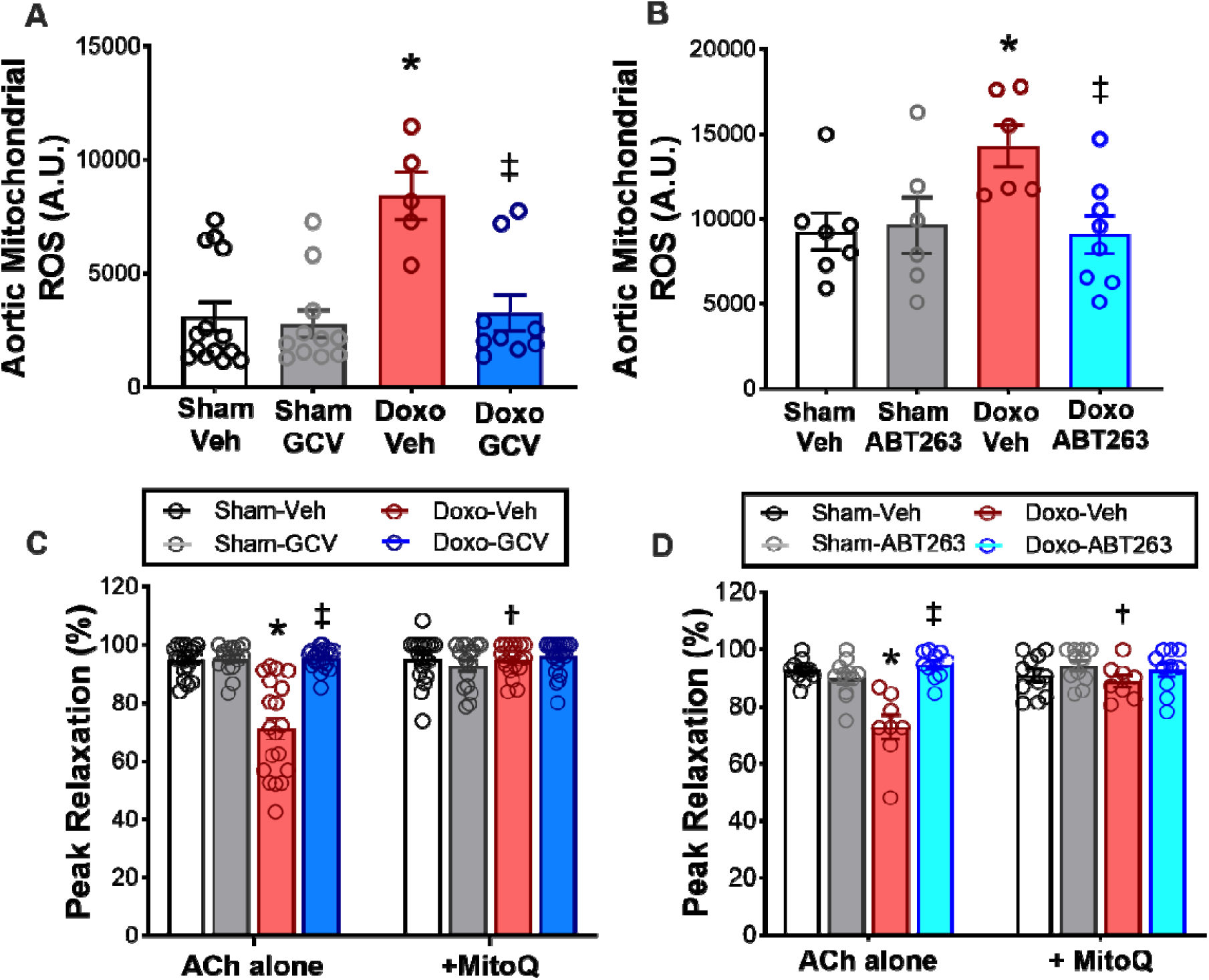
Doxo-induced cellular senescence promotes vascular mitochondrial related oxidative stress (mitoROS) which is prevented by senolytic treatment. Aortic mitochondrial reactive oxygen species (mitoROS) levels measured via electron paramagnetic resonance spectroscopy in study 1 **(A)** and 2 **(B)**. Carotid EDD in response to ACh in the presence/absence of mtROS scavenger, MitoQ in study 1 **(C)** and 2 **(D)**. All values are mean ± SEM. N=5-14/group in study 1 mitochondrial ROS production, N=6-10/ group in study 2 mitochondrial ROS production, N=13-25 in study 1 peak EDD with MitoQ, N=9-12 in study 2 peak EDD with MitoQ. *p<0.05 vs Sham-Veh, ‡p<0.05 vs Doxo-Veh.

Excessive mitoROS with Doxo could occur via reduced superoxide scavenging capacity due to lower abundance of the mitochondrial isoform of the superoxide dismutase (SOD) enzyme (SOD2) – a potent mitochondrial antioxidant in arteries^9,26^. Thus, we sought to determine if ABT263 influenced arterial SOD2 abundance following Doxo administration. We found that SOD2 abundance was not different (*P =* 0.51) in aorta lysates from Doxo-Veh mice relative to Sham-Veh animals and was not influenced by ABT263 treatment (**Figure S2B**), suggesting excessive mitoROS bioactivity with Doxo, and prevention with ABT263 treatment, is likely not influenced by a reduction in SOD2-mediated antioxidant defenses.

Lastly, we aimed to determine the causal role of the cellular senescence-mediated increase in mitoROS on conduit artery endothelial function. To accomplish this, we used a well-established bioassay for determining the direct role of excess mitoROS bioactivity in suppressing conduit artery endothelial function. Specifically, we assessed carotid artery EDD with and without prior incubation with the mito-targeted antioxidant MitoQ, as we have previously described^9,17^. MitoQ administration to the vessel perfusate restored peak EDD selectively in Doxo-Veh mice in study 1 and study 2 (*study 1*: *P* < 0.0001 with [95 ± 1%] vs. without [71 ± 4%] MitoQ; *study 2*: *P* = 0.001 with [89 ± 2%] vs. without [76 ± 5%] MitoQ) (**Figure 3C-D**). In studies 1 and 2, we did not observe an increase in peak EDD in the presence of MitoQ in the Doxo-GCV and Doxo-ABT263 groups, respectively (**Figure 3C-D**), suggesting that clearance of excess senescent cells following Doxo administration prevented the Doxo-induced effect of excessive mitoROS bioactivity on conduit artery endothelial function.

### Senolytic exposure prevents Doxo-induced endothelial dysfunction in the human microvasculature

We next aimed to translate our Doxo and senolytic findings in mice to the setting of the human microvasculature. To accomplish this, we leveraged a previously established human model of Doxo-induced microvascular endothelial dysfunction^18^ in which isolated human arterioles are exposed to Doxo (**Figure 4A**). In the present study, we exposed human adipose tissue arterioles to one of the four following experimental conditions: *1)* Control (sterile saline); *2)* ABT263 (2.5 μM); *3)* Doxo (100 nM); or *4)* Doxo + ABT263 and assessed both EDD and EID via flow-mediated dilation alone and with the smooth muscle dilator-papaverine, respectively. Arterioles incubated with Doxo had significantly lower peak EDD relative to control arterioles (Doxo, 32 ± 10% vs. Control, 94 ± 10%, *P* < 0.0001) which was fully prevented with ABT263 (Peak EDD: Doxo + ABT263, 82 ± 7%, *P* = 0.0001; *P* = 0.63 vs. Control; **Figure 4B**). Moreover, the influence of ABT263 was specific to Doxo-induced senescence, as evidenced by no difference between the Control and ABT263 groups (*P* = 0.96; **Figure 4B**). Incubation with papaverine (100 uM) abolished group differences (**Figure 4C**), thus suggesting that the observed differences with Doxo and ABT263 were specific to the endothelium. Overall, these data extend our preclinical findings in mice to the human microvasculature and further demonstrates that Doxo impairs endothelial function via excessive cellular senescence.

**Figure 4:**
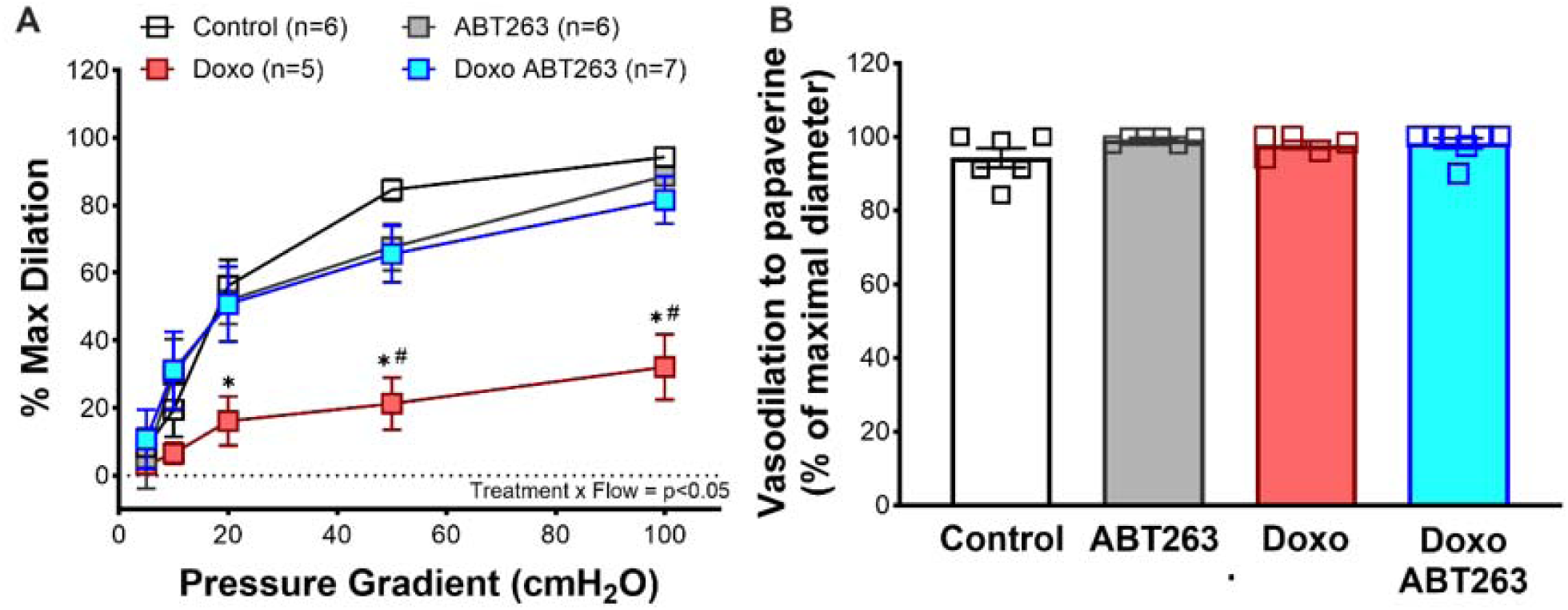
Senolytic exposure prevents doxorubicin (Doxo)-induced endothelial dysfunction in human arterioles. Arteriole endothelium-dependent dilation (EDD) response to increasing pressure gradient **(A)**. Vasodilation in response to the exogenous nitric oxide donor papaverine **(B)**. All values are mean ± SEM. N=5-7/group. *p<0.05 vs Sham-Veh, #p<0.05 vs Doxo-ABT263.

## DISCUSSION

In the present study, we used complementary experimental approaches, in male and female mice, to demonstrate the role of cellular senescence in mediating conduit artery endothelial dysfunction following Doxo administration. We also present translational evidence – in mice and isolated human arterioles – that supports the efficacy of targeting cellular senescence with senolytic therapy to mitigate endothelial dysfunction with Doxo. Collectively, these results provide proof-of-principle evidence that clearing excess senescent cells after Doxo administration prevents endothelial dysfunction and the use of senolytic therapy may be a viable option for accomplishing this goal.

Vascular endothelial dysfunction is a key antecedent to clinical CVD following Doxo chemotherapy treatment^3,6^. Clinical assessments of EDD in humans are independently predictive of CVD morbidity and mortality^27^. In this study, using carotid artery EDD, we found that cellular senescence mediates Doxo-induced endothelial dysfunction and that senolytic treatment following Doxo administration effectively prevents endothelial dysfunction. We have previously shown that Doxo-induced endothelial dysfunction is largely mediated by reduced NO bioavailability, primarily driven by excessive mitoROS^9^. However, the integrative mechanistic events influencing these responses remain incompletely understood. Here, we demonstrate that lower NO bioavailability following Doxo administration is mediated by excess cellular senescence and that senolytic therapy after Doxo can prevent this reduction. Furthermore, we show that Doxo-mediated cellular senescence induces excess vascular mitochondrial ROS bioactivity, which was directly implicated in Doxo-induced endothelial dysfunction, and that senolytic treatment prevents this effect. Notably, these changes in mitochondrial ROS appear to be independent of differences in antioxidant defenses, such as SOD2.

Doxo administration has also been shown to impair endothelial function in human microvessels^7,18^, though the underlying mechanisms were largely unknown. Here, we provide evidence that Doxo-induced cellular senescence plays an important role in mediating endothelial dysfunction in human microvessels. These observations offer novel translational evidence that cellular senescence is likely a key driver of Doxo-induced endothelial dysfunction in the human microcirculation and that senolytic therapy can prevent these effects.

Currently, there are multiple clinical trials investigating the efficacy of senolytic therapy for improving physiological function in cancer survivors. Notably, one trial (NCT04733534) is assessing whether reducing senescent cell burden with senolytic therapy (via administration of the chemotherapeutic agent dasatanib + the flavonoid quercetin or supplementation with the flavonoid fisetin) can improve frailty in adult survivors of childhood cancer. Additionally, a separate trial (NCT05595499) is evaluating the effects of fisetin supplementation on physical function in stage I–III breast cancer survivors. While these studies mark significant progress in the translation of senolytic therapy to clinical practice, there are no ongoing clinical trials specifically assessing the effects of senolytic therapy on endothelial function in cancer survivors, further demonstrating the need to establish preclinical proof-of-principle efficacy prior to clinical translation. Considering endothelial dysfunction plays a pivotal role in the development of CVD following cancer treatment, future research is needed to determine whether senolytic therapies can mitigate endothelial dysfunction in Doxo chemotherapy-treated cancer survivors.

In conclusion, this study provides key proof-of-principle evidence that cellular senescence plays critical mechanistic role in Doxo-induced endothelial dysfunction and that senolytic treatment following Doxo administration can prevent these adverse changes to endothelial function. Importantly, we demonstrate the role of cellular senescence in mediating additional mechanisms, including NO bioavailability and excessive mitochondrial oxidative stress, and how these events contribute to endothelial dysfunction induced by Doxo.

### Experimental Considerations and Future Directions

There are several important considerations in the present study. First, while we used healthy mice treated with Doxo to specifically assess the direct of chemotherapy on conduit artery endothelial function, this model does not account for the potential influence of the tumor microenvironment^28^. Since cancer itself can promote cellular senescence, this could contribute to endothelial dysfunction^29^; thus, future studies using cancer-bearing models would provide a more clinically relevant evaluation of Doxo-induced endothelial dysfunction. Second, although we demonstrate that excessive mitoROS contributes to Doxo-induced conduit artery endothelial dysfunction, we did not comprehensively assess other antioxidant defense systems beyond SOD2. Since senescent cells have altered redox balance and compromised antioxidant defenses^30^ exploring additional pathways, such as catalase or glutathione peroxidase, could offer further insight into how Doxo-induced cellular senescence more broadly influences redox balance and contributes to endothelial dysfunction. Third, while we translated our findings to human arterioles, the study did not assess systemic vascular function or clinical cardiovascular outcomes. Future studies are warranted to determine the safety of and establish the efficacy for senolytic therapy to improve endothelial function in Doxo chemotherapy-treated cancer survivors.

