## Supplementary material for "Cellular Senescence Mediates Doxorubicin Chemotherapy-Induced Vascular Endothelial Dysfunction: Translational Evidence of Prevention with Senolytic Treatment": Supplemetary materials

**SUPPLEMENTAL MATERIALS**

**Animals**

All male and female mice were housed in a conventional facility on a 12-hour light/dark cycle, given ad libitum access to an irradiated, fixed, and open standard rodent chow (Inotiv/Envigo 7917) and drinking water. All mice were euthanized by cardiac exsanguination while maintained under anesthesia (inhaled isoflurane) 2 to 4 weeks following the completion of the intervention periods (allowing for 1 week of recovery). After cardiac exsanguination, the carotid arteries were excised under a stereoscope, dissected free of connective tissue, and cannulated to fine glass canula within a pressure myograph system (DMT, Denmark) for assessment of vascular endothelial function. Investigators were blinded to the treatment group for data collection and biochemical analyses. All animal protocols were approved by the University of Colorado Boulder Institutional Animal Care and Use Committee (protocol no. 2618) and complied with the National Institutes of Health Guide for the Care and Use of Laboratory Animals.

**Study 1: Genetic-based clearance of senescent cells with GCV in p16-3MR mice**

p16-3MR mice were bred, weaned, and aged (to 4 months of age) in the animal care facility at the University of Colorado Boulder. At 4 months of age, male and female p16-3MR mice received either a single intraperitoneal injection of Sham (sterile saline) or Doxo (R&D Systems #2252/50 Minneapolis, MN) (10 mg/kg in Sham). One week later, mice either received the vehicle (Veh; saline) or ganciclovir (Sigma-Aldrich #G2536, St. Louis, MO) (GCV; 25 mg/kg/day in Veh) by intraperitoneal injection (IP) for 5 consecutive days, which is the standard approach for clearing senescent cells in this model, as we have described previously^1^. This equated to 4 groups/sex: Sham-Veh; Sham-GCV; Doxo-Veh; Doxo-GCV. For the primary outcomes of endothelial function, we studied n=13-25/sex/group. Throughout this intervention period, 5 mice died because of Doxo-related attrition, which resulted in a final sample size of Sham-Veh, n = 22; Sham-GCV, n = 13; DOXO-Veh, n = 23; and DOXO-GCV, n =25 .

**Study 2: Senolytic-based clearance of senescent cells with ABT263 in mice**

Male and female p16-3MR mice received a single intraperitoneal injection of Sham (sterile saline) or Doxo (10 mg/kg in Sham). One week later, mice either received the vehicle (Veh; 10% EtOH, 30% PEG400, 60% Phosal 50 PG) or the senolytic ABT263 (Selleckchem #S1001, Houston, TX) (50 mg/kg/day in Veh) by oral gavage on an intermittent one week on; two weeks off, one week on dosing paradigm, as we have previously described^1,2^. There were 4 groups/sex: Sham-Veh; Sham-ABT263; Doxo-Veh; Doxo-ABT263; n=10-12/group. Throughout this intervention period, 3 male mice died (2 due to Doxo-related attrition and one due to ABT263-related attrition), which resulted in a final sample size of Sham-Veh, n = 11; Sham-ABT263, n = 11; Doxo-Veh, n = 9; and Doxo-ABT263, n = 11 for our primary outcomes of vascular endothelial function.

**Study 3: *Ex vivo* human adipose tissue arteriole study – Influence of Doxo and ABT263**

Human arterioles were isolated from fresh surgical discard or consented tissue in accordance with the approved protocols by the Institutional Review Board of the Medical College of Wisconsin and Froedtert Hospital. Donor samples were received de-identified but with information about health status, co-morbidities and medication to select arterioles from healthy subjects defined as ≤1 risk factor for CVD. Subject demographics are presented in **Supplemental Table 3**. Healthy human arterioles were incubated with 100 nM Doxo (Sigma-Aldrich #D1515, St. Louis, MO) in PBS, 2.5 µM ABT263 in PBS or with a combination of both Doxo and ABT263 overnight (16-20h) like previously described^3^.

**Experimental procedures**

***Conduit artery endothelial function.*** After all the *in vivo* measurements were conducted following the completion of the intervention periods, mice were anesthetized with inhaled isoflurane and sacrificed via cardiac exsanguination and their carotids were extracted. Once excised, the carotids were dissected free of any perivascular adipose tissue and cannulated to glass micropipettes and placed under warm (37°C) physiological saline buffer in pressure myograph chambers (Danish Myograph Technology, A/S, Aarhus, Denmark). Once cannulated and pressurized to 50 mmHg intraluminal pressure, the carotids were allowed to rest for 45 minutes before conducting endothelium-dependent dilation (EDD) and endothelium-independent dilation (EID) in response to acetylcholine (ACh) and sodium nitroprusside (SNP) respectively, as described previously by our laboratory^1,2,4^. In brief, after the arteries were pre-constricted by phenylephrine (PE; 2 mM; Sigma-Aldrich, Cat# P6126), EDD was assessed by measuring the increases in the luminal diameter of the arteries in response to increasing concentration of ACh (1 X 10^-9^ to 1 X 10^-4^ M; Sigma-Aldrich, Cat# A6625). Following this, EID was assessed by measuring the luminal diameter in response to increased concentration of SNP; an exogenous NO donor (1 X 10^-10^ to 1 X 10^-4^ M; Sigma-Aldrich, Cat. No. 13755-38-9). After all the dose response measurement, the maximal luminal diameter of the vessel was measured after incubating it for 10-15 minutes in the Ca^2+^ free physiological saline buffer. All dose-response data are reported as percent changes in order to account for the baseline differences.

***NO mediated dilation.*** Carotid artery EDD was measured in presence of endothelial nitric oxide synthase inhibitor, *N*^G^-nitro-L-arginine methylester (L-NAME, 0.1mM, 30-min pre-incubation; Sigma-Aldrich, Cat# N5751) and NO mediated dilation was calculated by subtracting peak Ach dilation with peak L-NAME dilation as described by the equation below:

NO-mediated dilation (%) = Maximum dilation_Ach_ - Maximal dilation_Ach+L-NAME_

***Tonic mitoROS suppression of EDD*.** Carotid arteries were incubated for 60 minutes in the presence of a manganese superoxide dismutase mimetic mitoquinol mesylate (MitoQ, 1mMol/L). EDD was measured post incubation and mitochondrial ROS suppression was determined as the increase in EDD compared to Ach alone.

***Mitochondrial ROS production.*** Electron Paramagnetic Resonance (EPR), was used to measure mitochondrial-specific ROS production as previously described^4^. Using the mitochondrial-specific spin probe 1-hydroxy-4-[2-triphenylphosphonio)-acetamido]-2,2,6,6-tetramethylpiperidine (mitoTEMPO-H; Enzo Life Sciences), two 1-mm aortic rings were washed in warm physiological saline solution and incubated in Krebs/HEPES buffer, consisting of 99 mM NaCl, 4.7 mM KCl, 1.87 mM CaCl2, 1.2mM MgSO4, 25mM NaHCO3, 1.03mM KH2PO4, 20 mM Na-HEPES, 11.1 mM glucose, 0.1 mM diethy-lenetriaminepenta-acetic acid, 0.0035 mM sodium diethyldi-thiocarbamate, and Chelex (Sigma-Aldrich, Cat# C7901), containing 0.5 mM CMH at 37$^{\circ}$C for 60 minutes. MS300 Xband EPR spectrometer (Magnettech, Berlin, Germany) was used to analyze the samples under the following parameters: B0-Field, 3,350 G; sweep, 80 G; sweep time, 60 s; modulation, 3,000 mG; MW atten, 7 dB; gain, 5 x e^1.

***Sacrifice and tissue collection****.* Mice were sacrificed using a method approved under the American Veterinary Medical Association guidelines. Mice were anesthetized under inhaled anesthesia (open-drop method) and euthanized via cardiac exsanguination. The heart was removed, cleaned, and weighed. The aorta was excised and rinsed in physiological saline solution (PSS), cleared of perivascular adipose tissue and flash frozen for analysis later.

***Endothelial function in human adipose tissue arterioles.*** Human arterioles (100-300 μm in diameter) were dissected from de-identified surgical discard tissue kept in HEPES buffer, (275 mM NaCl, 20 mM HEPES acid, 12 mM glucose, 7.99 mM KCl, 4.9 mM MgSO_4_, 3.2 mM CaCl_2_·2H_2_O, 2.35 mM KH_2_PO_4_ and 0.07 mM EDTA). Following the removal of connective tissue, arterioles were cannulated on glass pipettes tips with matched impedance in an organ chamber containing Krebs buffer (123 mM NaCl, 19 mM NaHCO3, 11 mM glucose, 4,7 mM KCl, 2.5 mM CaCl_2_,1.2 mM MgSO_4_, 1.2 mM KH_2_PO_4_ and 0.026 mM EDTA). The buffer was supplied with 5% CO to ensure that the pH of 7.4 was maintained throughout the experiment. Following the cannulation, arterioles were subsequently pressured to 60 mmHg at 37°C for 30min. Endothelin-1 (max. 20nM) was utilized to pre-constricted vessels to 30-50% of their pressurized diameter and then subjected to five different flow gradients, ranging from 5 – 100 cmH2O, which equal shear rates of 5 to 25 dynes /cm^2^. EID at the end of max flow was assessed using 100 µM Papaverine.

***Aortic gene expression.*** mRNA gene expression was measured in segments of thoracic aorta following mechanical homogenization. RNA was extracted using the RNeasy mini kit (Qiagen). cDNA was synthesized using the iScript cDNA synthesis kit (Bio-Rad Laboratories, Hercules, CA). Transcripts of *Cdkn2a, Cdkn1a, Serpine1*, *Lmnb1, Cxcl1, Cxcl2, Ccl2, Plat1, Mmp3, Vegf, and Rage* were analyzed using a StepOnePlus Real-Time PCR System (Applied Biosystems, Waltham, MA) in 96-well plates and the Taqman OpenArray (Applied Biosystems) was used as a master mix, as described^2^. SimpleSeq DNA sequencing (Quintara Biosciences, Cambridge, MA) was used to validate PCR products. Primer sequences were as follows:

| **Gene** | **Species** | **Forward primer** | **Reverse primer** |
| --- | --- | --- | --- |
| *Cdkn2A* | Mouse | CCCAACGCCCCGAACT | GCAGAAGAGCTGCTACGTGAA |
| *Cdkn1A* | Mouse | TTGCCAGCAGAATAAAAGGTG | TTTGCTCCTGTGCGGAAC |
| *Serpine1* | Mouse | TGGAAGGGCAACATGACCAG | TCAGGCATGCCCAACTTCTC |
| *Lmnb1* | Mouse | GAGCCCCAAGAGCATCCAAT | CTGAGAAGGCTCTGCACTGT |
| *Cxcl1* | Mouse | CTGGGATTCACCTCAAGAACATC | CAGGGTCAAGGCAAGCCTC |
| *Cxcl2* | Mouse | CCTGGTTCAGAAAATCATCCA | CTTCCGTTGAGGGACAGC |
| *Ccl2* | Mouse | CACTCACCTGCTGCTACTCA | GCTTGGTGACAAAAACTACAGC |
| *Plat1* | Mouse | AAGCATGAGGCATCGTCTCC | ATGCATCGTGGAGGTCTTGG |
| *Mmp3* | Mouse | CTCGTGGTACCCACCAAGTC | CGCCAAAAGTGCCTGTCTTT |
| *Vegf* | Mouse | AAAAACGAAAGCGCAAGAAA | TTTCTCCGCTCTGAACAAGG |

**Aortic protein abundance**

***Immunoblotting.*** Protein abundance was measured in segments of thoracic aorta following mechanical homogenization in a bullet blender (Next Advance, Troy, NY) with zirconium oxide beads (3:1 1mm:0.5mm beads, Next Advance, Troy, NY) in a radioimmunoprecipitation assay lysis buffer supplemented with protease and phosphatase inhibitors [1 mMol/L sodium orthovanadate, 1X complete mini protease inhibitor cocktail tablet (Roche, Mannheim, Germany), 1 mMol/L phenylmethylsulfonyl fluoride, 1:100 Phosphatase Inhibitor Cocktail (Sigma-Aldrich, St. Louis, MO), 5 mMol/L sodium fluoride, and 5 mMol/L sodium pyrophosphate. Total protein content was quantified using a bicinchoninic acid assay (Thermo Fisher Scientific, Eugene, OR). Next, abundance of BAX (anti-rabbit; 1:50; Cell Signaling; Danvers, MA Cat. No. 2772), BCL-2 (anti-goat; 1:50; R&D Systems, Minneapolis, MN Cat. No. AF810); Manganese superoxide dismutase (SOD2) (antigoat; 1:50; R&D Systems, cat no. AF3419) were determined by loading 0.4 μg/μL of aortic protein per capillary in a 25-lane (capillary) automated Western blot quantitative analyzer (JESS, ProteinSimple, San Jose, CA), according to the manufacturer’s guidelines, as described previously^1,6^, following the validation of these antibodies in test aorta lysates. Anti-goat and anti-rabbit secondary antibodies were provided by the manufacturer and used according to the manufacturer’s guidelines. A grayscale analysis of the band intensities was then performed to quantify protein abundance using Compass software (ProteinSimple), with target proteins expressed relative to total protein.

***Immunofluorescence.*** Immunofluorescence assays were performed to visualize the subcellular localization of target proteins; nitrotyrosine. At the time of sacrifice, ~1mm sections of thoracic aorta were excised and frozen in OCT (Tissue-Tek^®^ O.C.T.) compound, as described above^1,^ . Later in time, 7 μm sections (Leica CM1520) were plated on poly-L-lysine-coated microscope slides, fixed in 4% paraformaldehyde, washed with PBS, and permeabilized (0.1% Triton X-100). Slides were then incubated with anti-nitrotyrosine primary antibody (1:50; Cell Signaling Technology, Danvers, MA; Cat# 9691) overnight, washed with PBS, and incubated with a species-specific fluorescent secondary antibody (AlexaFluor 647; Invitrogen, Waltham, MA) for 60 minutes. Slides were washed, stained with DAPI (1:1000; Invitrogen, Waltham, MA; Cat# D1306) for 5 minutes, and cured overnight with ProLong Gold mounting media (Invitrogen, Waltham, MA; Cat# P36980). The slides were then imaged using EVOS m7000 (ThermoFisher, Waltham, MA; Cat# AMF7000) fluorescence microscope under identical conditions and analyzed using Invitrogen Celleste 5.0 Image Analysis Software. Images for were obtained at 10x magnification. The abundance of nitrotyrosine was determined as the average intensity (A.U.) of the positively stained area across N = 6-8 samples/animal.

**Statistical Analysis**

Statistical analyses were conducted in Prism, version 10 (GraphPad Software, Inc. La Jolla, CA, USA). Data were first assessed for outliers (ROUT method, *Q* = 1%) and normality (Shapiro-Wilk normality test, *P* > 0.05) within groups. Differences in EDD and EID, both of which are terminal measurements obtained upon sacrifice, were assessed using a two-way mixed ANOVA (group x dose). Differences across animal groups in morphological and artery characteristics, NO-mediated dilation, mitoROS bioactivity, and qPCR were assessed using one-way ANOVA. Differences in nitrotyrosine expression were analyzed via unpaired t-test. When significant main effects were detected, pairwise comparisons were made using the Holm-Sidak post hoc test. Significance was set to α = 0.05. Unless otherwise noted, data are presented as mean ± SEM.

**Supplementary figures**

**Supplementary Table 1** Characteristics of mice in the doxorubicin (DOXO) ABT-263 study

|  | Sham  Veh | Sham ABT263 | DOXO  Veh | DOXO  ABT263 |
| --- | --- | --- | --- | --- |
| *n* | 12  *3 female*  *9 male* | 11  *6 female*  *5 male* | 10  *4 female*  *6 male* | 11  *5 female*  *6 male* |
| Body weight, g | 28.1 ± 1.9 | 27.6 ± 1.6 | 26.2 ± 0.9 | 24.4 ± 1.8 |
| Heart mass, mg | 142.8 ± 8.3 | 140.5 ± 5.9 | 149.3 ± 6.8 | 137.0 ± 9.0 |
| Left ventricle mass, mg | 76.5 ± 5.2 | 71.1 ± 6.2 | 69.5 ± 4.3 | 65.7 ± 4.9 |
| Quadriceps mass, mg | 302.3 ± 9.3 | 304.5 ± 10.3 | 308.0 ± 9.0 | 296.1 ± 15.0 |
| Visceral adipose mass, mg | 727.7 ± 17.1 | 592.5 ± 13.8 | 301.9 ± 5.7* | 280.7 ± 8.6* |
| Spleen mass, mg | 65.4 ± 3.9 | 68.1 ± 3.4 | 76.5 ± 5.9 | 67.3 ± 5.5 |
| Carotid artery |  |  |  |  |
| Resting diameter, μM | 466 ± 9 | 460 ± 7 | 454 ± 6 | 459 ± 6 |
| Maximal diameter, μM | 505 ± 8 | 503 ± 7 | 506 ± 8 | 502 ± 6 |

Data are mean ± SEM. **P*<0.05 vs. Sham-Vehicle (Veh).

**Supplementary Table 2** Subject Characteristics

| **Parameter** | **Total** |
| --- | --- |
| **N** (female/male) | 10 (10/0) |
| **Age**, years (mean ± SEM) | 41.3 ± 3.2 |
| **Ethnicity/race** |  |
| Asian, n (%) | 0 (0%) |
| Black, n (%) | 3 (30%) |
| Hispanic, n (%) | 0 (0%) |
| Non-Hispanic white, n (%) | 6 (60%) |
| Did not report, n (%) | 1 (10%) |
| **Adipose Tissue**  Subcutaneous, n (%) 10 (100%)  **Cardiovascular risk factors** | |
| Hypertension, n (%) | 0 (0%) |
| Hyperlipidemia, n (%) | 0 (0%) |
| Coronary artery disease, n (%) | 0 (0%) |
| Tobacco use, n (%) | 0 (0%) |
| Congestive heart failure, n (%) | 0 (0%) |
| Other | 0 (0%) |
| COVID (8/2022) | 1 (10%) |
| Breast Cancer  **Medication** | 1 (10%) |
| Beta blocker, n (%) | 1 (10%) |
| Calcium blocker, n (%) | 0 (0%) |
| ACE inhibtiors, n (%) | 0 (0%) |
| Nitrates, n (%) | 0 (0%) |

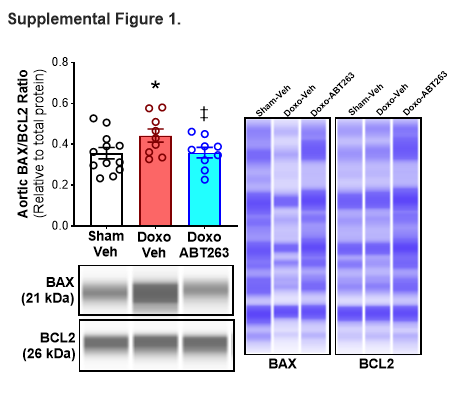

**Supplementary figure 1: Doxo administration increases aortic Bax:Bcls2 ratio and senolytic treatment with ABT263 prevents it.** Bax:Bcl2 ratio for aortas in study 2 along with representative images. All values are mean ± SEM. N=9-12/group *p<0.05 vs Sham-Veh, ‡p<0.05 vs Doxo-Veh.

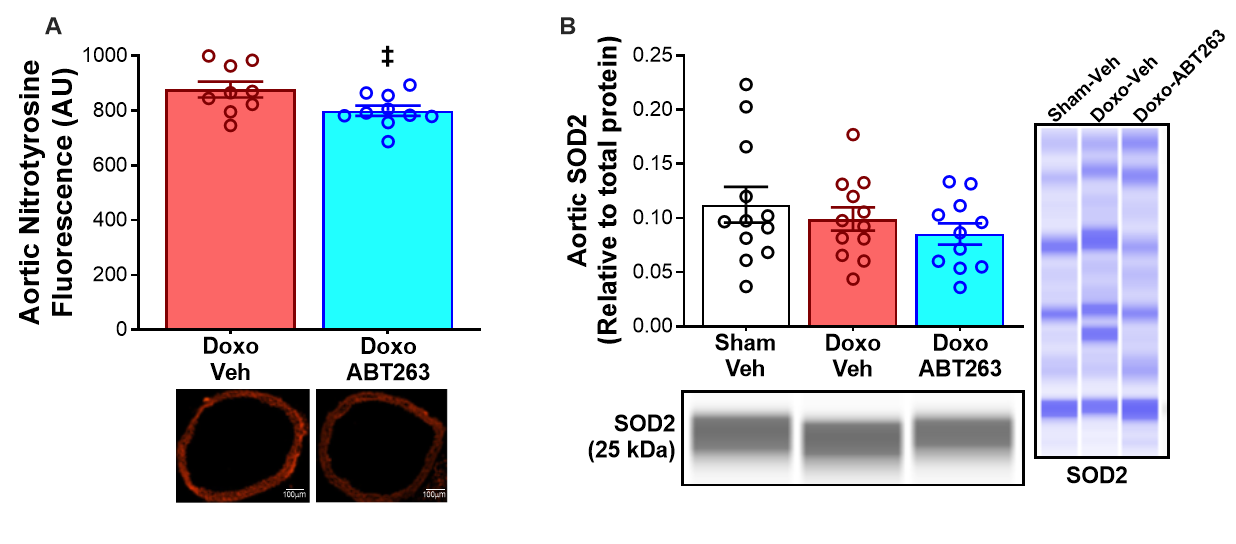

**Supplementary figure 2: Doxo administration increases aortic nitrotyrosine levels and senolytic treatment prevents it independent of SOD2 levels**. Immuflurescence (IF) levels of aortic nitrotyrosine levels in study 2 **(A)**. Aortic manganese superoxide dismutase (SOD2) protein expression in study 2 **(B)**. All values are mean ± SEM. N=6-12/group *p<0.05 vs Sham-Veh, ‡p<0.05 vs Doxo-Veh.
